# Simple and effective serum-free medium for sustained expansion of bovine satellite cells for cell cultured meat

**DOI:** 10.1101/2021.05.28.446057

**Authors:** Andrew J. Stout, Addison B. Mirliani, Eugene C. White, John S.K. Yuen, David L. Kaplan

## Abstract

Cell-cultured meat offers the potential for a more sustainable, ethical, resilient, and healthy food system. However, research and development has been hindered by the lack of suitable serum-free media that enable the robust expansion of relevant cells (e.g., muscle satellite cells) over multiple passages. Recently, a low-cost serum-free media (B8) was described for induced pluripotent stem cells. Here, we adapt this media for bovine satellite cells and show that the addition of a single component, recombinant albumin, renders B8 suitable for the long-term expansion of cells without sacrificing myogenicity. We show that this new media (Beefy-9) maintains robust cell growth over the entire culture period tested (seven passages) with an average growth rate of 39 hours per population doubling. Along with demonstrated efficacy for bovine cells, this work provides a promising starting-point for developing serum-free media for cultures from other meat-relevant species. Ultimately, this work offers a promising foundation for escaping the reliance on serum in cultured meat research, thereby accelerating the field.

## Introduction

Cell-cultured meat is an emerging technology which currently offers both promising possibilities and significant scientific challenges. The promise of cultured meat lies in its potential to address environmental, ethical, and human health issues that plague intensive animal agriculture^1^. For instance, limited life-cycle analyses suggest that cultured meat could require >90% less land and >75% less water than conventional beef, while contributing >75% fewer greenhouse gas emissions, >95% less eutrophication, and >90% less particulate matter formation^2,3^. At the same time, cultured meat could improve animal welfare, food security, and human health outcomes^4–6^. The challenges that face the successful technological transition of cultured meat to the marketplace stem from the need for production systems that are low-cost, scalable, food-safe, and free of animal-derived inputs^4,7,8^. Here, cell culture media is a particularly problematic hurdle for several reasons. First, cell culture media comprises the majority (>99%) of the cost of current production systems^7–9^. Second, the culture of meat-relevant cells (e.g., bovine satellite cells) has traditionally relied on fetal bovine serum (FBS), a notoriously expensive, unsustainable, and inconsistent component, which is inherently antithetical to the aims of cultured meat^4^. Finally, when serum-free media for satellite cells have been explored, they are either complex^10^, ineffective compared to serum-containing media^11^, reliant on proprietary or animal-derived additives^10–12^, or contain additives (e.g., synthetic steroids) that could cause regulatory concern^12^. Further, no serum-free media has been validated for the sustained expansion of muscle satellite cells over multiple passages^10–12^. As such, serum-free media remains one of the most pressing limitations for the field.

Recently a simple, low-cost, serum-free medium was described for expanding human iPSCs^13^. This media (B8) is based on the commonly used Essential 8 medium and contains a simple mixture of basal media (glucose, amino acids, vitamins, salts, and fatty acids), minerals, and proteins, all of which can be found in animal tissue; as such, it is likely to be considered food safe. Lab-scale B8 production was demonstrated at a cost of ∼$16/L when using in-house production of growth factors. It is likely that this cost could be further reduced with process scale-up and optimization. In contrast, media containing 20% serum or commercially available serum-free media that have been explored for bovine satellite cell expansion cost ∼$200-500/L ^11,13^. Because of its simple and cost-effective nature, B8 offers a promising starting point for establishing an effective serum-free media that could enhance the relevance and pace of cultured meat research.

Here, we validate a simple, B8-inspired serum-free media, termed “Beefy-9”, for culturing primary bovine satellite cells (BSCs). First, we show that supplementation with just one animal-free component (recombinant human albumin, expressed in rice) makes Beefy-9 media effective for BSC expansion *in vitro*, and that short-term growth rates are comparable to those obtained in media containing 20% FBS. Next, we establish a protocol for passaging BSCs in Beefy-9 and show that passaged cells maintain their myogenicity in serum-free conditions. Third, we demonstrate cost-saving opportunities in Beefy-9 formulation by lowering growth-factor concentrations without a significant reduction in proliferation. Finally, we validate the long-term expansion and sustained myogenicity of BSCs in serum-free conditions. Together, these findings offer a simple and valuable resource for cultured meat researchers, and a promising foundation for further media optimization to enable low-cost, scalable, and sustainable cultured meat production.

## Results

### B8 media can lower serum requirements for BSCs during short-term growth

To test B8’s capacity to replace serum containing media, short term BSC growth (3 and 4 days) was analyzed in mixtures of BSC-GM combined with homemade B8 or supplier provided HiDef-B8 (Fig. 1 & Fig. S1). These timepoints were selected because cells in fast-growing conditions reached confluency by day 5 (data not shown). The results showed that B8 media mixed with BSC-GM significantly improved growth compared to BSC-GM alone over four days (Fig. 1A), and that this benefit remained with as much as a 62.5% reduction in FBS (62.5% B8 media). Additionally, an 87.5% reduction in serum did not significantly reduce cell growth over four days. On the other hand, while B8 media alone encouraged cell growth over three days (Fig. 1A & B), there was no change between days 3 and 4, indicating that growth did not continue into day 4. These results indicated that B8 was capable of reducing serum requirements in BSCs, but could not completely eliminate these requirements.

**Figure 1:**
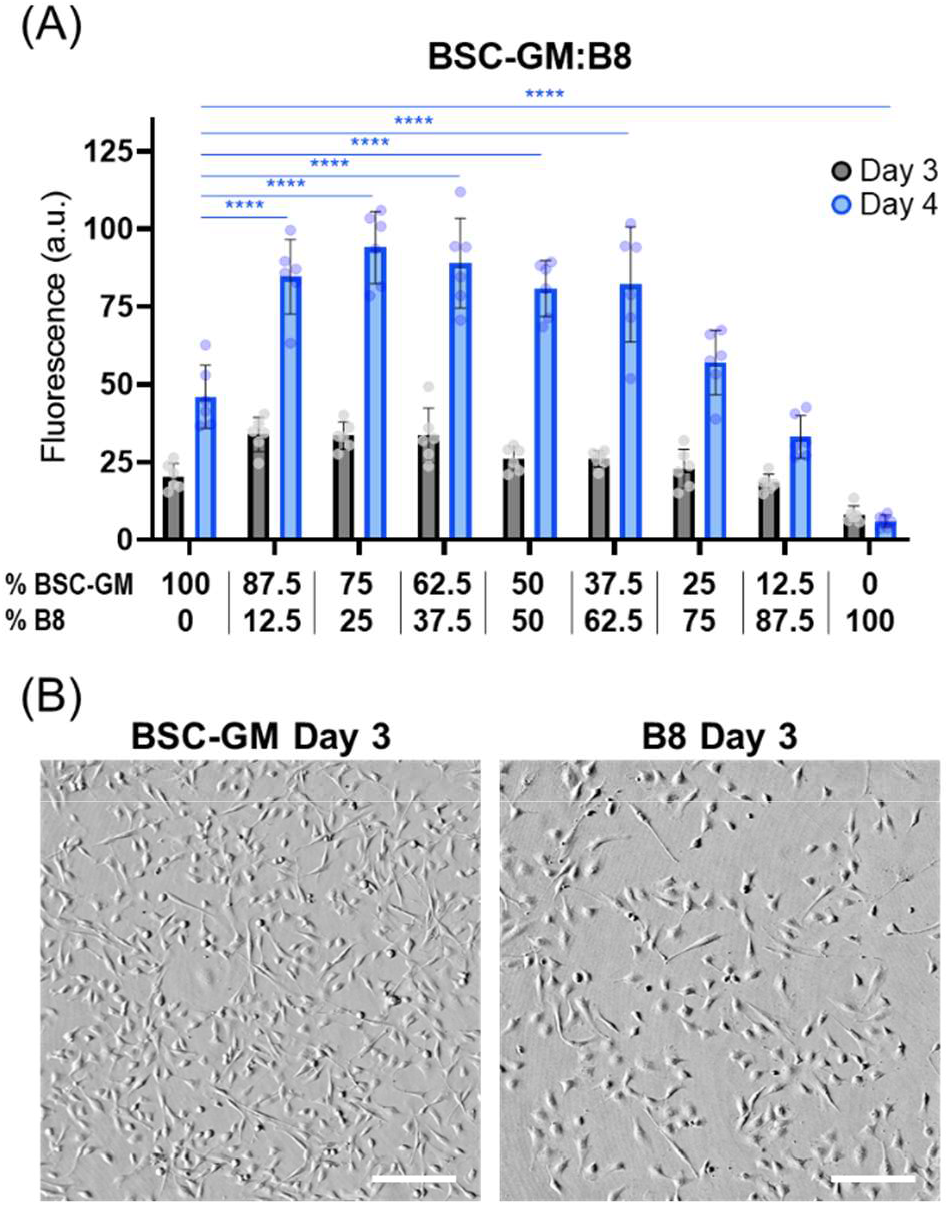
Short-term growth in BSC-GM mixed with B8. **(A)** BSC proliferation over 3 & 4 days in mixtures of BSC-GM (20% FBS) and B8 media. At the four-day timepoint, mixtures of up to 62.5% B8 significantly improved growth compared to BSC-GM alone, and mixtures of up to 87.5% B8 did not significantly reduce growth (p=0.27). B8 alone showed a significant reduction in growth over four days compared with BSC-GM alone, and showed stagnating growth after three days. n=6 distinct samples; statistical significance was calculated by one-way ANOVA on day 4 data comparing all samples with BSC-GM controls, and is indicated by asterisks, in which p < 0.05 (*), p <0.01 (**), p < 0.001 (***), and p < 0.0001 (****). **(B)** Brightfield images of BSCs grown for three days in BSC-GM or B8 media. Images show that cell morphology was consistent in serum-containing or serum-free conditions. Cell confluency in the images is qualitatively consistent with growth analysis in part (A). Scale bars 200 μm.

### B8 media supplementation improves cell proliferation

To overcome the deficiencies of B8 media alone, numerous supplements were tested in a range of concentrations (full list in Table S1), and growth was again analyzed over four days. Six of the factors significantly improved BSC proliferation compared to B8 media alone (Fig. 2A & Fig. S2). These were interleukin-6 (IL-6), curcumin, recombinant human albumin (rAlbumin), Platelet-derived growth factor (PDGF-BB), linoleic acid, and oleic acid. Of these, rAlbumin was particularly effective, imparting a ∼4-fold improvement in growth compared with plain B8. In contrast, other supplements resulted in at best only a ∼50% improvement compared with plain B8. To test whether combinations of these supplements could offer synergistic benefits to cell growth, optimal concentrations of the above factors (with the exception of PDGF-BB, which was determined to be insufficiently effective for the substantial cost) were combined and tested (Fig. 2B). Here, rAlbumin (800 μg/mL) was the driving factor in all significant improvements. While a combination of IL-6 (0.01 ng/mL) and rAlbumin offered slightly improved growth compared with rAbumin alone, this difference was not statistically significant, and in the interest of maximizing media simplicity and minimizing media cost, an augmented B8 media with nine components and designed for bovine muscle culture (thus termed Beefy-9) was established by supplementing with 800 μg/mL rAlbumin alone. This media was capable of maintaining short-term growth comparable to serum-containing BSC-GM and maintaining cell morphology *in vitro* (Fig. 2B & C).

**Figure 2:**
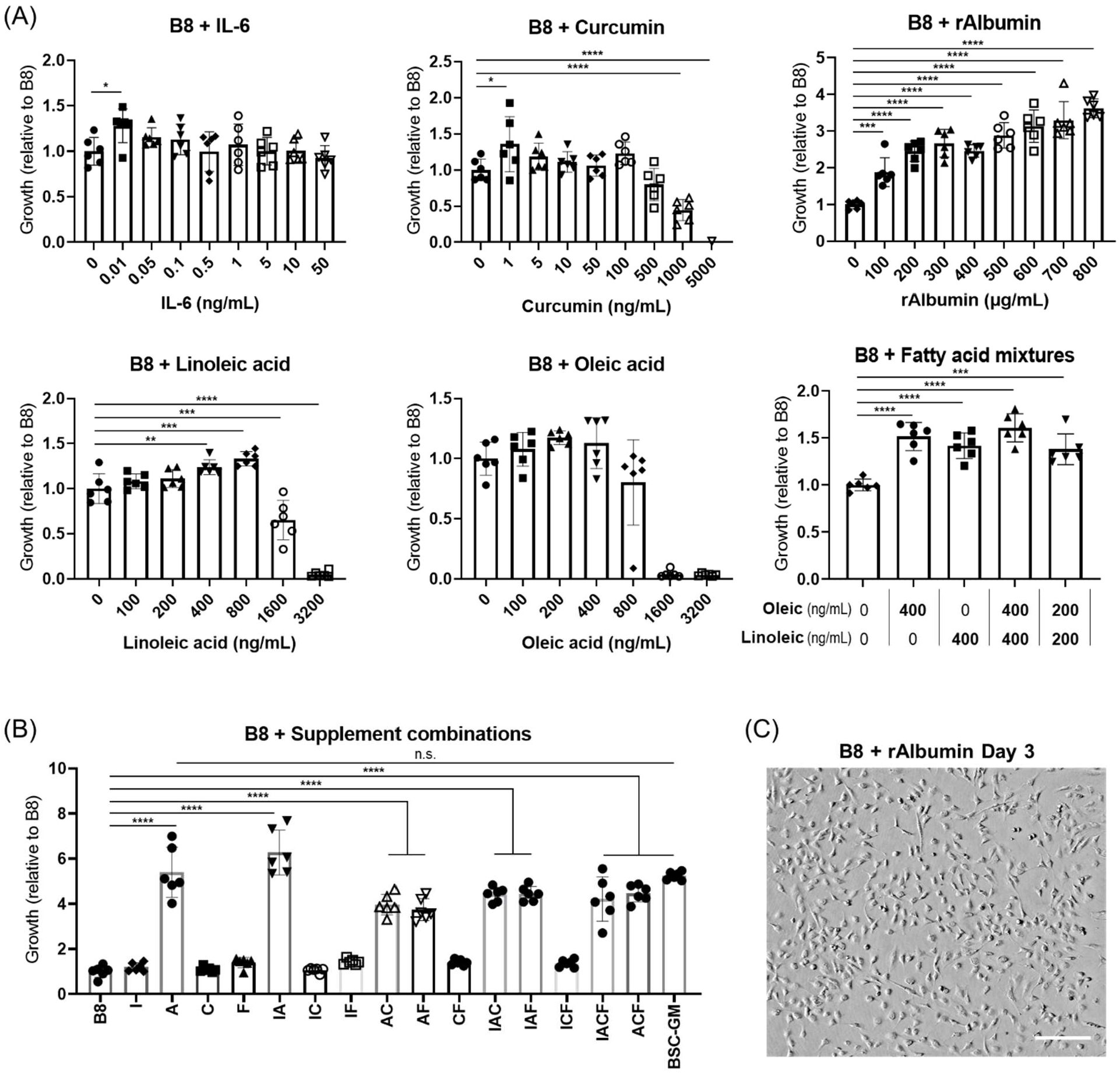
Short-term growth in supplemented B8 media. **(A)** BSC proliferation over 4 days B8 supplemented with Interleukin 6 (IL-6), curcumin, recombinant albumin (rAlbumin), linoleic acid, oleic acid, or mixtures of linoleic and oleic acid. Values are given relative to B8. n=6 distinct samples; statistical significance was calculated by one-way ANOVA with multiple comparisons between supplemented samples and B8 controls, and is indicated by asterisks, in which p < 0.05 (*), p <0.01 (**), p < 0.001 (***), and p < 0.0001 (****). **(B)** BSC proliferation over 4 days in B8 supplemented with combinations of factors, in which: B8 = B8; I = IL-6 (.01 ng/mL); A = rAlbumin (800 μg/mL); C = Curcumin (1 ng/mL); F= linoleic acid (400 ng/mL) and Oleic acid (400 ng/mL); and BSC-GM = serum-containing growth media. n=6 distinct samples; statistical significance was calculated by one-way ANOVA with multiple comparisons between all samples, and is indicated by asterisks, in which p < 0.05 (*), p <0.01 (**), p < 0.001 (***), and p < 0.0001 (****). Statistically significant differences between B8 and other samples are shown, as is the insignificance between rAlbumin supplemented B8 and BSC-GM (p>0.9990). **(C)** Brightfield imaging of BSCs grown for four days in B8 supplemented with rAlbumin (800 μg/mL) media shows that cell morphology was maintained in B8 with albumin supplementation compared with images in Figure 1. Scale bar 200 μm.

### Passaging in Beefy-9 media

While short-term growth experiments were useful in establishing the benefits of rAlbumin supplementation for tailoring B8 media to BSC short term culture needs, long-term culture and passaging are essential for robust cell expansion required for cultured meat. However, seeding cells into Beefy-9 media directly after passaging initially proved ineffective, as cells did not re-adhere to tissue culture plastic. We hypothesized two possible explanations for this result. The first was that the coating used (0.25 μg/cm^2^ of laminin-511) was insufficient to enable cell adhesion in the absence of other adhesion factors present in serum (e.g., vitronectin and fibronectin)^14^. The second was that the high concentration of albumin was outcompeting laminin for adsorption to the tissue-culture plastic, thereby further hindering cell adhesion^15,16^. To overcome these possible limitations, we explored passaging BSCs in the absence of albumin and adding albumin one day after plating, in order to allow cells to adhere to the flasks, as well as passaging cells with various concentrations of different adhesive proteins. Because one priority of this work was media and cell culture workflow simplicity, we focused on recombinant adhesive proteins that were relatively low-cost and which have been demonstrated without pre-coating [e.g., laminin-511 fragment (iMatrix-511) and truncated vitronectin (Vtn-N)]^17,18^. We also explored Poly-D-Lysine (PDL) coatings (which can be purchased commercially on pre-coated plates) to augment cell attachment with or without adhesive peptides^18^. The results indicated that delaying the addition of rAlbumin was necessary to allow cell adhesion and growth, as is coating with a cell adhesive peptide such as iMatrix-511 (Fig. 3A). However, when comparing various cell adhesive peptides, the results indicated that iMatrix-511 was sub-optimal compared with Vtn-N (Fig. 3B). Specifically, 1.5 μg/cm^2^ of Vtn-N showed superior cell adhesion (day 1) and growth (day 4) than PDL alone, laminin alone, PDL+laminin, or a lower concentration of Vtn-N with or without PDL. Once a suitable passaging method was determined, short-term growth curves were again performed with Beefy-9 supplemented with various growth factors in order to rule out the possible confounding effect that adsorbed serum proteins might have had on short term growth curves performed previously (in which cells were seeded in the presence of serum). No significant effect of these factors was found (Fig. S3), reaffirming that supplementation with rAlbumin alone was optimal over multiple passages.

**Figure 3:**
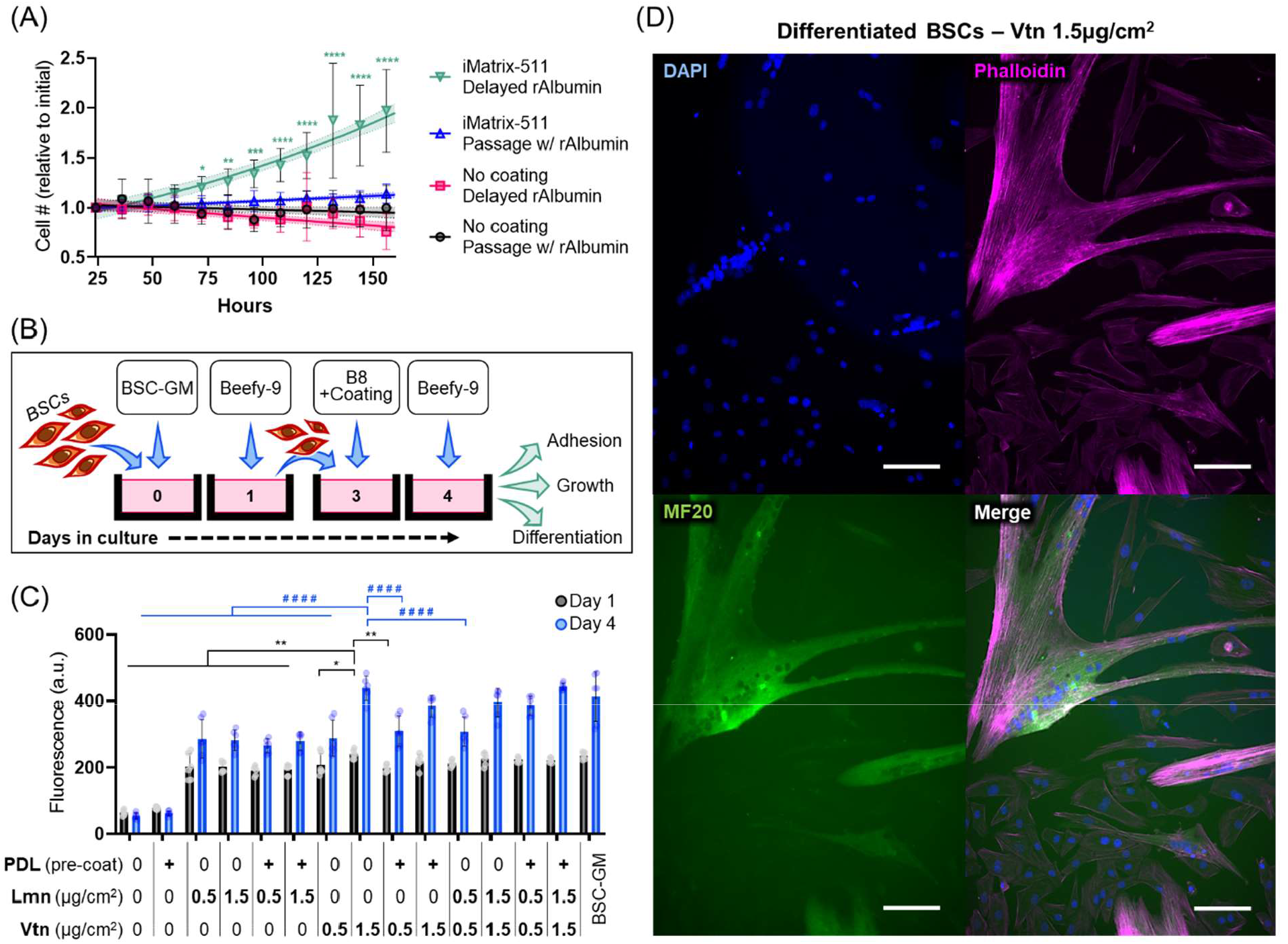
Passaging in Beefy-9. **(A)** Growth analysis of BSCs passaged in B8/Beefy-9 media. Results showed that cells needed to be passaged in the absence of supplemental Albumin (“Delayed rAlbumin”), and that a coating (e.g., iMatrix-511 laminin) was required for adhesion and growth. Specifically, cells without any coating (“No coating”) were unable to grow, as were cells with iMatrix-511 coating that were passaged in the presence of albumin (“iMatrix-511; Passage w/ rAlbumin”). In contrast, cells that were passaged onto iMatrix-511 coated flasks and allowed to adhere overnight before the addition of albumin (“Delayed rAlbumin”) showed exponential growth. n=9 image fields of view; statistical significance calculated by two-way ANOVA with multiple comparisons between conditions, and significant difference between “iMatrix-511; Delayed rAlbumin” and all other conditions are indicated by asterisks, in which p < 0.05 (*), p <0.01 (**), p < 0.001 (***), and p < 0.0001 (****). A 95% confidence interval calculated via nonlinear regression (least squares regression; exponential (Malthusian) growth). **(B)** Schematic of B8/Beefy-9 passaging system used. BSCs were plated in BSC-GM on day 0. On day 1, cells were rinsed with DPBS and media was changed to Beefy-9. At 70% confluency (day 3), cells were passaged using TrypLE and plated in B8 (no albumin) along with various adhesive peptides. One day after passaging, media was again changed to Beefy-9, and cells were proliferated and analyzed for adhesion, growth, and myogenicity. **(C)** PrestoBlue adhesion and growth analysis of BSCs plated with various animal-free coatings. Truncated vitronectin (Vtn-N) at 1.5 μg/cm^2^ shows superior cell attachment and growth compared to iMatrix-511 laminin (Lmn) or Poly-D-Lysine (PDL). n=3 distinct samples and 2 technical replicates; statistical significance was calculated by one-way ANOVA performed separately for day 1 or day 4 with multiple comparisons between Vtn-N 1.5 μg/cm^2^ and all other samples. and is indicated by asterisks (day 1) or hashes (day 4), in which p< p < 0.05 (*, #), p <0.01 (**, ##), p < 0.001 (***, ###), and p < 0.0001 (****, ####). **(D)** Immunofluorescence staining for nuclei (DAPI, blue), actin (Phalloidin, magenta), and Myosin Heavy Chain (MF20, green) in BSCs passaged in Beefy-9 media with 1.5 μg/cm^2^ Vtn-N and delayed rAlbumin. Cells were proliferated to confluency and differentiated in a previously described serum-free differentiation medium. Scale bars 50 μm.

### Serum-free differentiation following expansion in Beefy-9 media

After establishing delayed rAlbumin and 1.5 μg/cm^2^ of Vtn-N as suitable parameters for multiple-passage culture of BSCs in Beefy-9, the myogenicity of expanded cells was determined. Specifically, Beefy-9-passaged cells (P2) were expanded to confluency in Beefy-9 and differentiated for 5 days in a previously published serum-free differentiation medium^10^. Differentiated cells showed the formation of multinucleated myotubes which stained positive for the myogenic marker myosin heavy chain (MHC) (Fig. 3C). The results validated that myogenicity of cells cultured and passaged in Beefy-9 media was maintained. Together, these results demonstrate a fully animal-component-free culture system for proliferating, passaging, and differentiating BSCs (Supplementary Video 1).

### Cost-reducing strategies for Beefy-9 media

We next explored simple cost reduction strategies for Beefy-9 by lowering the concentration of FGF-2, which is a major cost contributor to B8 and Beefy-9 formulations at the baseline concentration of 40 ng/mL. Growth was analyzed over four days as before in B8 and Beefy-9 media with FGF-2 concentrations ranging from 0-80 ng/mL. The results showed that for B8 and Beefy-9, FGF-2 could be lowered to 5 ng/mL and 1.25 ng/mL, respectively, without significantly affecting growth (Fig. 4A). In contrast, cell growth and morphology were significantly affected by the complete removal of FGF-2 from the media (Fig. 4A & B). The results indicated that substantial reduction in FGF-2 is possible to lower the cost of Beefy-9 without negatively impacting short-term growth rates.

**Figure 4:**
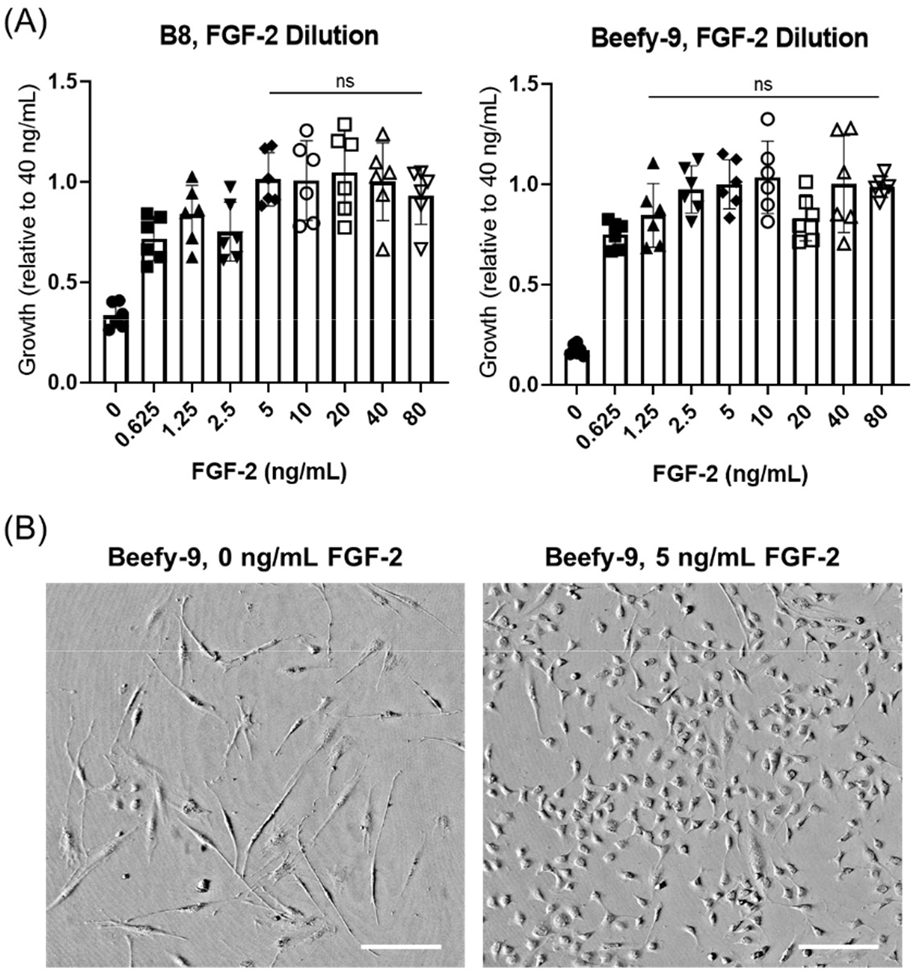
Short-term growth in B8 & Beefy-9 with reduced FGF-2. **(A)** BSC proliferation over 4 days in B8 or Beefy-9 with various concentrations of FGF-2. FGF-2 could be reduced to 5 ng/mL or 1.25 ng/mL in B8 or Beefy-9, respectively, without significantly impacting cell growth over four days. n=6 distinct samples; statistical significance was calculated by one-way ANOVA with multiple comparisons between various FGF-2 concentrations and 40 ng/mL control conditions, and is indicated by asterisks, in which p < 0.05 (*), p <0.01 (**), p < 0.001 (***), and p < 0.0001 (****). **(B)** Brightfield images of BSCs grown for three days in Beefy-9 media with 0 or 5 ng/mL FGF-2. Images show that the complete removal of FGF-2 from Beefy-9 significantly affects cell morphology, whereas a reduction to 5 ng/mL did not affect morphology as compared with images in Figures 1 & 2. Scale bars 200 μm.

### Long-term culture in Beef-9 media

The last step after validating Beefy-9 for short-term growth and establishing an appropriate passaging protocol was to validate long-term expansion of BSCs in Beefy-9. BSCs were seeded as before and fed with either serum-containing BSC-GM, B8, Beefy-9 with high FGF-2 (40 ng/mL), or Beefy-9 with low FGF-2 (5 ng/mL). The low FGF-2 concentration was conservatively selected as the concentration that did not significantly affect short-term growth in either B8 or Beefy-9. Cells were cultured and passaged as described (Fig. 3B) for seven passages over 28 days, with cell counts used to determine cumulative cell doublings over the four-week period. While BSC-GM was still the optimal media over a long growth period, Beefy-9 with high or low FGF-2 content showed significant improvements over B8 media without the addition of rAlbumin (Fig. 5A). Indeed, BSCs in B8 alone ceased proliferating after three passages (4.4 doublings), while BSCs in Beefy-9 continued to expand exponentially over at least seven passages (18.2 and 17.2 doublings for 40 ng/mL FGF-2 and 5 ng/mL FGF-2, respectively).

**Figure 5:**
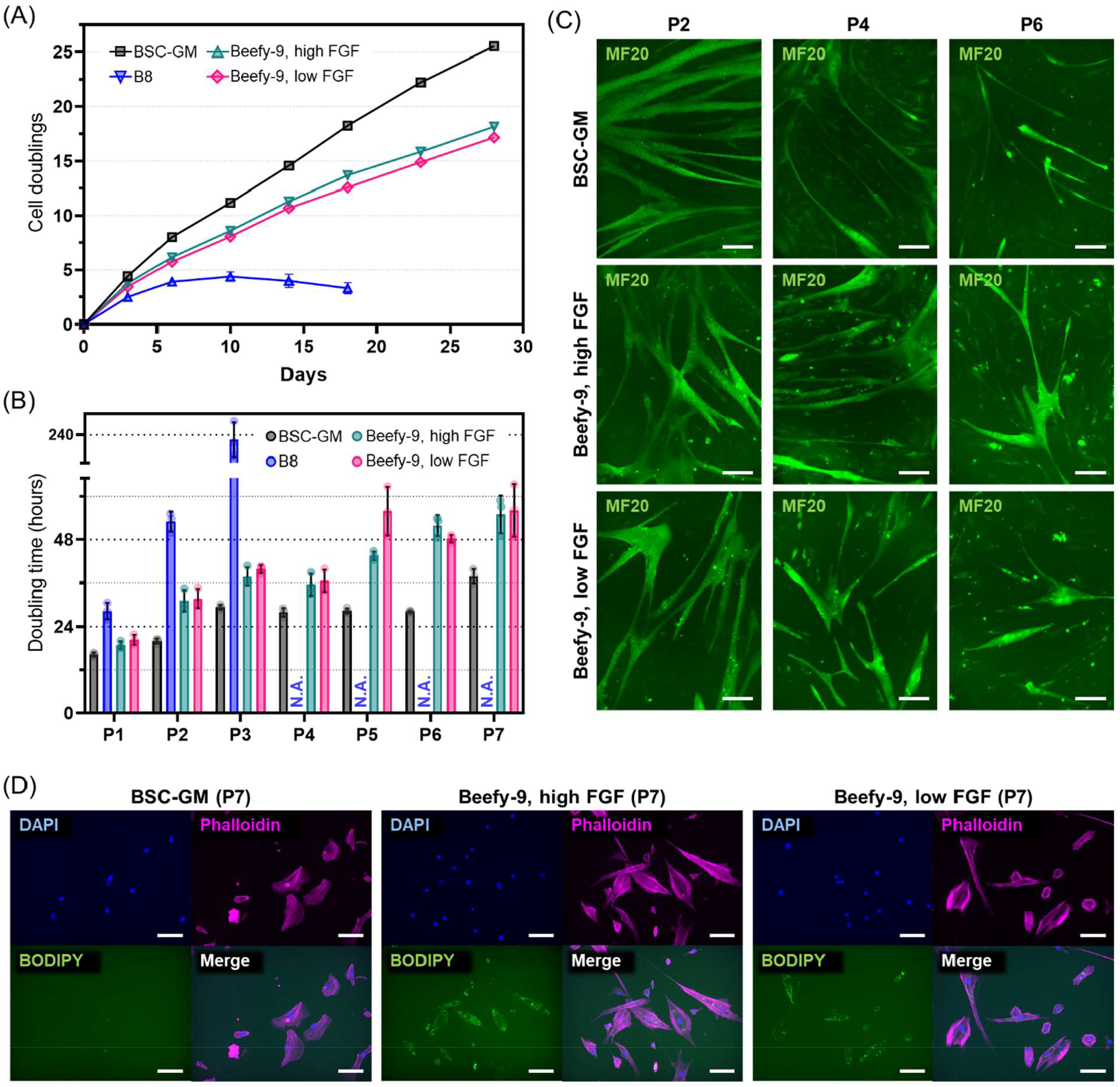
Long-term culture. **(A)** Cell doublings over multiple passages of BSCs cultured in BSC-GM, B8, Beefy-9, high FGF (40 ng/mL FGF-2), or Beefy-9, low FGF (5 ng/mL FGF-2). Results show that the addition of albumin significantly improved cell growth in B8, although not to the degree of serum, over four weeks. Reducing Beefy-9 FGF-2 to 5 ng/mL decreased cell doublings compared to 40 ng/mL, although this difference was less substantial (17.2 doublings at 28 days for low FGF-2 vs 18.2 doublings for high FGF-2). **(B)** Doubling times were calculated over long-term cell culture and compared between media types. An increase in doubling time at higher passages was found, particularly for Beefy-9 with high or low FGF-2. Notably, however, doubling times remained <48 hours for the first five passages (∼13 doublings) in Beefy-9. **(C)** Immunofluorescence staining for Myosin Heavy Chain (MF20, green) in BSCs at passages 2, 4 and 6 during long-term culture experiments in BSC-GM, Beefy-9, and low-FGF Beefy-9. After reaching confluency, cells were differentiated in a previously described serum-free differentiation medium^10^ for four days (P2 and P4) or six days (P6) and stained for MHC. Images revealed that myogenicity of BSCs was maintained throughout long term culture in all media. Scale bars 200 μm. **(C)** BODIPY staining of lipid accumulation in P7 cells cultured in BSC-GM, Beefy-9, high FGF, and Beefy-9, low FGF. Cells were passaged, allowed to adhere overnight, and stained for nuclei (DAPI, blue), actin (Phalloidin, magenta), and lipids (BODIPY, green). Images revealed increased lipid accumulation in BSCs cultured over seven passages in serum-free media compared to serum-containing media. Scale bars 50 μm.

Converting growth data to doubling times revealed a consistent increase in doubling time over seven passages in Beefy-9 media, with a slower increase in doubling time in BSC-GM (Fig. 5B); however, doubling times remained below 48 hours for Beef-9 media for the first five passages (13.7 doublings) and under 56 hours for all seven passages (18.2 doublings). The average doubling time over seven passages in Beefy-9 with high or low FGF-2 concentrations were ∼39 and ∼41 hours, respectively. These are higher than the doubling time of ∼17 hours that has been reported for satellite cells *in vivo*, but within a range expected based on previous reports of BSCs cultured *in vitro*^19–21^. Together, these results indicated that the Beefy-9 media with both high and low FGF-2 concentrations were effective for the long-term expansion of bovine satellite cells, but that further optimization is needed to improve growth rates over multiple passages.

### BSC phenotype and myogenicity over long-term culture

Throughout long-term culture, the myogenicity of BSCs was verified in serum-containing and serum-free conditions. Cells from passages 2, 4, and 6 were cultured to confluency, differentiated in serum-free differentiation medium, and stained for Myosin Heavy Chain (MHC) as before. The formation of MHC-positive multinucleated myotubes in BSC-GM and Beefy-9 formulations over six passages was shown (Fig. 5C). Interestingly, differentiation appeared improved in BSCs cultured in Beefy-9 media compared with BSC-GM over multiple passages, both in terms of myotube size and density, and in terms of quantitative fusion index (Fig. 5C & Fig. S4). This result could be because cells cultured in BSC-GM have undergone more doublings than cells cultured in Beefy-9 at the points of analysis, or because non-myogenic cells (e.g., fibroblast-like cells) overgrow myogenic satellite cells more rapidly in serum-containing media. While myogenicity was maintained throughout expansion in Beefy-9 media, it was noted that cells cultured in serum-free conditions also appeared to accumulate lipid droplets over long-term culture, while cells cultured in BSC-GM did not (Fig. 5D). This aberrant lipid accumulation could be due to insulin resistance in cells as a result of the relatively high concentration of insulin in B8 & Beefy-9, and could therefore point towards a possible media optimization strategy by adjusting insulin levels^22,23^. Alternately, lipid accumulation could suggest that BSCs are thrust towards an adipogenic phenotype in Beefy-9 media^11^, though the sustained myogenicity of BSCs in serum-free conditions affirms the capacity of these media to maintain relevant satellite cell function for cultured meat.

### Media cost analysis

Once the efficacy of Beefy-9 was demonstrated, we performed a simple cost analysis to understand how this media might be implemented by research groups currently relying on serum-containing media (e.g., 20% FBS + 1 ng/mL FGF-2, as used in this study) for cultured meat research (Fig. 6). Price comparisons revealed that even using purchased growth factors and without bulk ordering (as in this study), Beefy-9 media cost substantially less than serum-containing media ($217/L vs. $290/L, respectively). At the reduced 5 ng/mL of FGF-2, the price of Beefy-9 drops further to $189/L). Further fold price decreases could easily be achieved by increasing the scale of culture media component orders, and by using powdered basal media. Specifically, the costs for Beefy-9 with high or low FGF-2 concentrations dropped to $74/L and $46/L, respectively, when components are ordered in bulk (component sourcing given in Table S3). In this study, Beefy-9 prices were dominated by rAlbumin, basal media, FGF-2 (at high concentrations), and insulin. If bulk ordering of store-bought components is used, the price was dominated by FGF-2 (at high concentrations), rAlbumin, and insulin, with basal media offering significantly less impact. While Beefy-9 is easy to produce in-house, further ease-of-use can be achieved by purchasing HiDef-B8 media and simply adding rAlbumin to prepare HiDef-Beefy-9; however, at current prices this results in a significant increase in cost.

**Figure 6:**
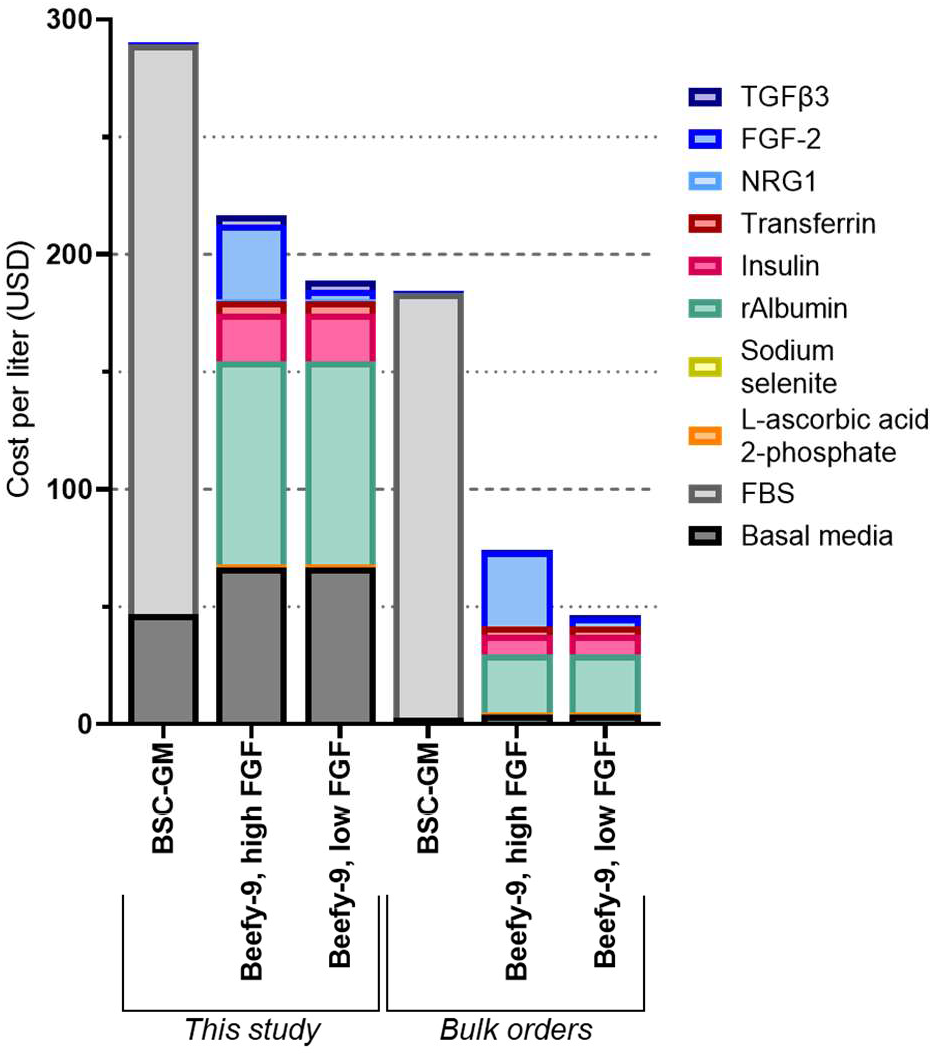
Cost analysis. Media costs for BSC-GM, Beefy-9, high FGF, and Beefy-9, low FGF were assessed for reagents bought at small scale (those used in this study), and reagents if bought in bulk from available suppliers. For both small-scale and bulk purchasing, Beefy-9 (high and low FGF) are both more economical than BSC-GM. When purchased in bulk and using low FGF concentrations, Beefy-9 can be prepared for less than $50/L. For both small-scale and bulk purchasing, rAlbumin is a major cost-driver for Beefy-9 media, as is FGF-2 when used at 40 ng/mL.

## Discussion

Since its emergence, cultured meat research and development has been stymied by a lack of suitable serum-free media for muscle stem cell expansion. This deficiency has led the field to rely on FBS for most research, thereby hindering the relevance of findings. This is particularly the case if we look forward to serum-free production processes. Developing serum-free media for relevant cell types (e.g., muscle and fat) and relevant species (e.g., bovine, porcine, galline, etc.) is therefore essential to accelerate research in the field. These media should at a minimum be affordable, comprised of food-safe components, reliable, and easy to use (e.g., not overly complex) in order to promote their adoption in widespread research efforts and towards scaled production processes. In this work, we describe Beefy-9 media as a promising candidate for simple serum-free culture of bovine satellite cells. We validated: 1) the efficacy of this medium in promoting BSC proliferation, 2) the ability of this media to enable long-term expansion when used in combination with rAlbumin-free B8 and truncated vitronectin, and 3) the maintenance of satellite cell myogenicity when expanded in Beefy-9.

The significant impact of rAlbumin on Beefy-9’s efficacy for BSCs is noteworthy, particularly considering that the addition of albumin did not show nearly as profound of an effect on iPSC growth when B8 was originally developed^13^. Albumin has many roles in cell culture media, and indeed comprises about 60% of the total proteins in serum. For an in-depth summary of these roles, the reader is directed to several thorough reviews^24,25^. Briefly, though, albumin acts in culture media to bind, carry, and stabilize compounds such as fatty acids, metal ions, signaling molecules, amino acids, and other factors. As such, it is a potent and multifaceted antioxidant which can sequester these species from redox or other degrading reactions, increasing the half-life, availability, and solubility of beneficial factors to cells while reducing the accumulation of harmful byproducts. It is possible that these regulating effects impart an advantage to Beefy-9 over B8 alone, as the latter was not incapable of promoting cell division in BSCs over the short term, only of fostering robust and prolonged expansion. Interestingly, albumin has also been suggested as a potential protectant against cell stresses due to sparging bubbles in various bioreactors, and so Beefy-9 could offer additional benefits in these scaled-up bioprocesses^24^.

Cost analysis suggests that Beefy-9 media should be easy to implement and affordable for academic labs that are currently using serum-containing (e.g., 20% FBS) media for cultured meat research. It should be noted, however, that significant cost reduction of serum-free media is still needed to make cultured meat production economically viable from an industrial perspective. Here, reducing the cost of albumin, growth factors and basal media (e.g., through the use of plant or algal hydrolysates) will be essential^26,27^. Additionally, co-culture of meat-relevant cells with nutrient- or growth factor-producing cells could offer valuable cost-saving opportunities^28^. When considering Beefy-9, it is clear that recombinant proteins are the main drivers of cost. As such, further research should explore cost-reduction of these recombinant proteins, or opportunities for substituting or eliminating these factors. Additionally, it should be noted that in the absence of cell-adhesion proteins found in FBS, a significant increase in the concentration of tissue-vessel coating was required to achieve adequate BSC adhesion and growth during serum-free culture. This factor is often overlooked when discussing the cost of large-scale cell culture; however, the present study relied on 1.5 μg/cm^2^ of Vtn-N, which adds ∼$0.18/cm^2^, or $31.75 per T-175 flask (a standard vessel used in our lab, suitable for ∼5 doublings at standard seeding and passaging densities). Opportunities to reduce the costs associated with cell adhesion include adapting or engineering cells to suspension culture, reducing the cost of recombinant adhesion protein production, or exploring low-cost alternatives to recombinant production^4,29,30^.

While this work presents a promising resource for researchers and a useful foundation for further media development, it is clear that further media optimization for both cost and long-term efficacy is warranted. Specifically, the present work relies on one-factor-at-a-time exploration of media components to find suitable supplements to tailor B8 for bovine muscle stem cells. As media components are intimately entangled in their effects on cell biology, it is therefore unlikely that Beefy-9 is optimal as-is. Indeed, it is possible that media components which appeared inconsequential in this work (e.g., hepatocyte growth factor or ethanolamine) would offer advantages in other permutations of media not tested in the development of Beefy-9. Computational approaches to media development are better suited for solving such multi-factorial problems, and should therefore be leveraged to further optimize Beefy-9 and other serum-free media for cultured meat applications^27^. Here, efforts can focus on adipose as well as muscle tissue, for instance through the use of free fatty acid addition to Beefy-9 to induce the transdifferentiation of BSCs into lipid-accumulating cells^31^. Additionally, further work is needed to overcome the slower long-term growth and aberrant lipid accumulation that is seen in BSCs cultured over seven passages in Beefy-9. Promising options for this include the use of spontaneously or genetically immortalized stem cells, which could improve long-term outcomes in Beefy-9^32,33^.

The present work offers a simple, affordable, and effective Beefy-9 serum-free media for improving cultured meat research. However, the cost and efficacy of Beefy-9 will require further optimization for industrial scale production of cultured meats that aim to reach price-parity with conventional meats. Future work to address these needs could focus on both engineering efforts (e.g., increased recombinant growth factor productivity, production of species-specific recombinant proteins, optimized media formulations, and cell line engineering) and scientific discoveries (e.g., novel protein analogues or alternatives, or insights into cell signaling pathways) in order to drive the cost of production down. Here, cultured meat development is likely to provide collateral benefit to biomedical research, such as tissue engineering for volumetric muscle loss or cell-based biopharmaceutical production. Ultimately, we expect that sustained efforts towards serum-free media development will continually lower costs and improve scalability of cultured meat over time, bringing products closer to market viability and cultured meat’s possible benefits closer to reality.

## Materials and methods

### Primary bovine satellite cell isolation and maintenance

Primary bovine satellite cells (BSCs) were isolated with methods previously used in our group^6^. Briefly, ∼0.5 g of muscle was excised from the semitendinosus of a 14-day-old Simmental calf at the Tufts Cummings School of Veterinary Medicine according to approved methods (IACUC protocol #G2018-36). Muscle tissue was minced into a paste and digested in 0.2% collagenase II (Worthington Biochemical #LS004176, Lakewood, NJ, USA; 275 U/mg) for 45 minutes with regular trituration. Digestion was halted with BSC growth media (BSC-GM) comprised of DMEM+Glutamax (ThermoFisher #10566024, Waltham, MA, USA) supplemented with 20% fetal bovine serum (FBS; ThermoFisher #26140079), 1 ng/mL human FGF-2 (ThermoFisher #68-8785-63), and 1% Primocin (Invivogen #ant-pm-1, San Diego, CA, USA), and cells were filtered and plated at a density of 100,000 cells/cm^2^ onto uncoated tissue-culture flasks. After 24 hours of incubation at 37°C with 5% CO_2_, the plated suspensions (containing satellite cells) were transferred to flasks coated with 1μg/cm^2^ mouse laminin (Sigma #CC095, St. Louis, MO, USA), which were left untouched for three days before growth media was changed, and cells were cultured using standard practices on tissue-culture plastic coated with 0.25 ug/cm^2^ iMatrix recombinant laminin-511 (Iwai North America #N892021, San Carlos, CA, USA). After two weeks of culture, Primocin in growth media was replaced with 1% antibiotic-antimycotic (ThermoFisher #1540062). For regular cell maintenance, cells were cultured at 37°C in 5% CO_2_ to a maximum of 70% confluence, counted using an NC-200 automated cell counter (Chemometec, Allerod, Denmark), and either passaged using 0.25% trypsin-EDTA (ThermoFisher #25200056) or frozen in FBS with 10% Dimethyl sulfoxide (DMSO, Sigma #D2650). The identity and myogenicity of BSCs used in this study was previously validated by our group^6^.

### Short-term growth analysis

Homemade B8 medium was prepared using store-bought components and a previously described formulation and method of preparation (Table S1)^34^. Additionally, HiDef-B8 medium aliquots were generously provided by Defined Bioscience (Defined Bioscience # LSS-201, San Diego, CA, USA) and added DMEM/F12 with 1% antibiotic/antimycotic. Short term BSC growth (3 and 4 days) was analyzed for mixtures of serum-containing and serum-free media, as well as for pure B8 media with reduced growth factor concentrations and/or with the addition of various media supplements (Table S1). Briefly, BSCs were thawed (passage number < 2) and plated in BSC-GM on 96-well tissue-culture plastic plates for each timepoint at a density of 2,500 cells/cm^2^ with 0.25 ug/cm^2^ iMatrix recombinant laminin-511. After 24 hours, BSC-GM was removed, cells were washed 1x with DPBS, and new media (e.g., B8 +/-supplementation) was added. A list of supplements and concentrations can be found in Table S1. Media was changed on day 3, and on days 3 and 4 cells were imaged, media was aspirated from appropriate plates, and plates were frozen at -80°C. Once all timepoints were frozen, cell number was analyzed using a FluoReporter dsDNA quantitation kit (ThermoFisher #F2962) according to the recommended protocol and with fluorescence readings performed on a Synergy H1 microplate reader (BioTek Instruments, Winooski, VT, USA) using excitation and emission filters centered at 360 and 490 nm, respectively. Cell number at 3 and 4 days was analyzed relative to pure B8 or HiDef-B8 media.

### Passaging in Beefy-9 media

To test various passaging conditions using B8 + rAlbumin (Beefy-9), BSCs were plated in BSC-GM onto T-75 culture flasks at a density of 2,500 cells/cm^2^ with 0.25 ug/cm^2^ iMatrix recombinant laminin-511. After 24 hours, BSC-GM was removed, cells were washed 1x with DPBS, and Beefy-9 media was added. Cells were cultured to 70% confluency, harvested with TrypLE Express (ThermoFisher #12604021), centrifuged at 300g, and resuspended in B8 or Beefy-9 media with or without iMatrix laminin-511. Cells were seeded at 5,000 cells/cm^2^ (0.25 ug/cm^2^ iMatrix laminin) onto a 12-well plate, and growth was analyzed with a live cell monitoring system (Olympus Provi CM20, Tokyo, Japan). After 24 hours, media was aspirated, and all cells were fed with Beefy-9 media. Cell growth was compared over seven days in order to determine the effects of seeding +/-rAlbumin and +/-iMatrix laminin.

To test the effect of different coatings, 48-well plates were prepared with or without pre-coating with poly-D-lysine (Sigma #P1024-10MG) according to the manufacturer’s instructions. Cells were then cultured and harvested as above, centrifuged at 300g, and resuspended in B8 media with varying concentrations if iMatrix laminin and/or truncated recombinant human vitronectin (ThermoFisher #A14700). Cells were seeded at 5,000 cells/cm^2^. After 24 hours, media was aspirated, and cells were rinsed 1x with DPBS. Cells were then fed with Beefy-9 media with 10% PrestoBlue reagent (ThermoFisher #A13262) and incubated at 37°C. After 2.5 hours, PrestoBlue media was moved to a 96-well plate and read with a Synergy H1 microplate reader using excitation and emission filters centered at 560 and 590 nm, respectively. Cell culture media was then replenished with Beefy-9 and PrestoBlue analysis was repeated on Day 4.

### Long-term growth analysis

For generating long-term growth curves, BSCs were thawed and plated (P1) onto 6-well culture plates (triplicate wells) in BSC-GM with 0.25 ug/cm^2^ iMatrix laminin-511. After allowing cells to adhere overnight, media was removed, cells were washed 1x with DPBS, and either BSC-GM, B8, Beefy-9 (40 ng/mL FGF-2) or Beefy-9 (5 ng/mL FGF-2) were added to cells. Upon reaching ∼70% confluency, cells were rinsed with DPBS, harvested with TrypLE Express, and counted using an NC-200 automated cell counter (duplicate counts for each well). Cells were then pelleted at 300g, resuspended in BSC-GM or B8 media, re-counted, and seeded onto new 6-well plates at 2,500 cells/cm^2^ with 0.25 ug/cm^2^ iMatrix laminin-511 (cells in BSC-GM) or 1.5 ug/cm^2^ recombinant vitronectin (cells in B8). After allowing cells to adhere overnight, media was replaced with appropriate media (BSC-GM, B8, Beefy-9 (40 ng/mL FGF-2) or Beefy-9 (5 ng/mL FGF-2). This process was repeated over 28 days and seven passages. Throughout culture, cells were fed every two days. When seeding cells for passages 2, 4, 6, and 7, additional wells were seeded for staining for myosin heavy chain (P2, P4 and P6) or lipid accumulation (P7).

### Immunocytochemistry

To analyze the effect of B8 and Beefy-9 media on cellular phenotype, BSCs cultured in serum-free media were stained for Paired-box 7 (Pax7), a marker of satellite cell identity. Proliferative cells in BSC-GM, B8, and Beefy-9 media were fixed with 4% paraformaldehyde (ThermoFisher #AAJ61899AK) for 30 minutes, washed in PBS, permeabilized for 15 minutes using 0.5% Triton-X (Sigma # T8787) in PBS, blocked for 45 minutes using 5% goat serum (ThermoFisher #16210064) in PBS with 0.05% sodium azide (Sigma #S2002), and washed with PBS containing 0.1% Tween-20 (Sigma #P1379). Primary Pax7 antibodies (ThermoFisher #PA5-68506) were diluted 1:100 in blocking solution containing 1:100 Phalloidin 594 (ThermoFisher #A12381), added to cells, and incubated overnight at 4°C. Cells were then washed with PBS + Tween-20, incubated with secondary antibodies for Pax7 (ThermoFisher #A-11008, 1:500) for 1 hour at room temperature, washed with PBS + tween-20, and mounted with Fluoroshield mounting medium with DAPI (Abcam #ab104139, Cambridge, UK) before imaging. Imaging was performed via fluorescence microscopy (KEYENCE, BZ-X700, Osaka, Japan).

To verify myogenicity of BSCs in various media, cells were cultured to confluency in serum-containing or serum-free conditions, and media was changed to a previously described serum-free differentiation media consisting of Neurobasal (Invitrogen #21103049, Carlsbad, CA, USA) and L15 (Invitrogen #11415064) basal media (1:1) supplemented with 1% antibiotic/antimycotic, 10ng/mL insulin-like growth factor 1 (IGF-1; Shenandoah Biotechnology #100-34AF-100UG, Warminster, PA, USA) and 10 ng/mL epidermal growth factor (EGF; Shenandoah Biotechnology #100-26-500UG). Following 5-7 days of differentiation, cells were fixed, stained, and imaged as previously described, using primary antibodies for MHC (Developmental studies hybridoma bank #MF-20, Iowa City, IA, USA), phalloidin 594 (1:100), appropriate secondary antibodies (ThermoFisher #A-11001, 1:1000), and Fluoroshield mounting medium with DAPI.

### Statistical analysis

Statistical analysis was performed with GraphPad Prism 9.0 software (San Diego, CA, USA). Short-term cell growth analyses were performed via one-way or two-way ANOVA, as appropriate, with multiple comparisons performed with the Tukey’s HSD post-hoc test. Regression analysis (Fig. 3A) was performed via nonlinear regression (least squares; exponential (Malthusian) growth) and is shown alongside 95% confidence interval. Doubling time (Fig. 5B) was determined through nonlinear regression (least squares; exponential (Malthusian) growth) for each biological replicate (well) of each passage, using technical replicates (duplicate counts) in generating nonlinear regressions. P values <0.05 were treated as significant. Unless otherwise stated, errors are given as ± standard deviation.

## Author contributions

**Andrew Stout**: Conceptualization, Methodology, Investigation, Formal Analysis, Visualization, Writing-original draft preparation. **Addison Mirliani:** Investigation, Writing-original draft preparation. **John Yuen:** Conceptualization, Writing-reviewing and editing. **Eugene White:** Methodology, Resources. **David Kaplan:** Conceptualization, Resources, Writing-reviewing and editing, Supervision, Funding acquisition.

## Acknowledgements

We thank New Harvest for their support of this work. We also thank Dr. Steven Rees and Defined Bioscience for their scientific input and generous donation of HiDef-B8 media. We thank Natalie Rubio for her assistance with the satellite cell isolations, and Scott Brundage for his help obtaining bovine tissues. Thanks also go to Natalie Rubio, Kyle Fish, Ning Xiang, Michael Saad, and Sophie Letcher for their scientific input.

---

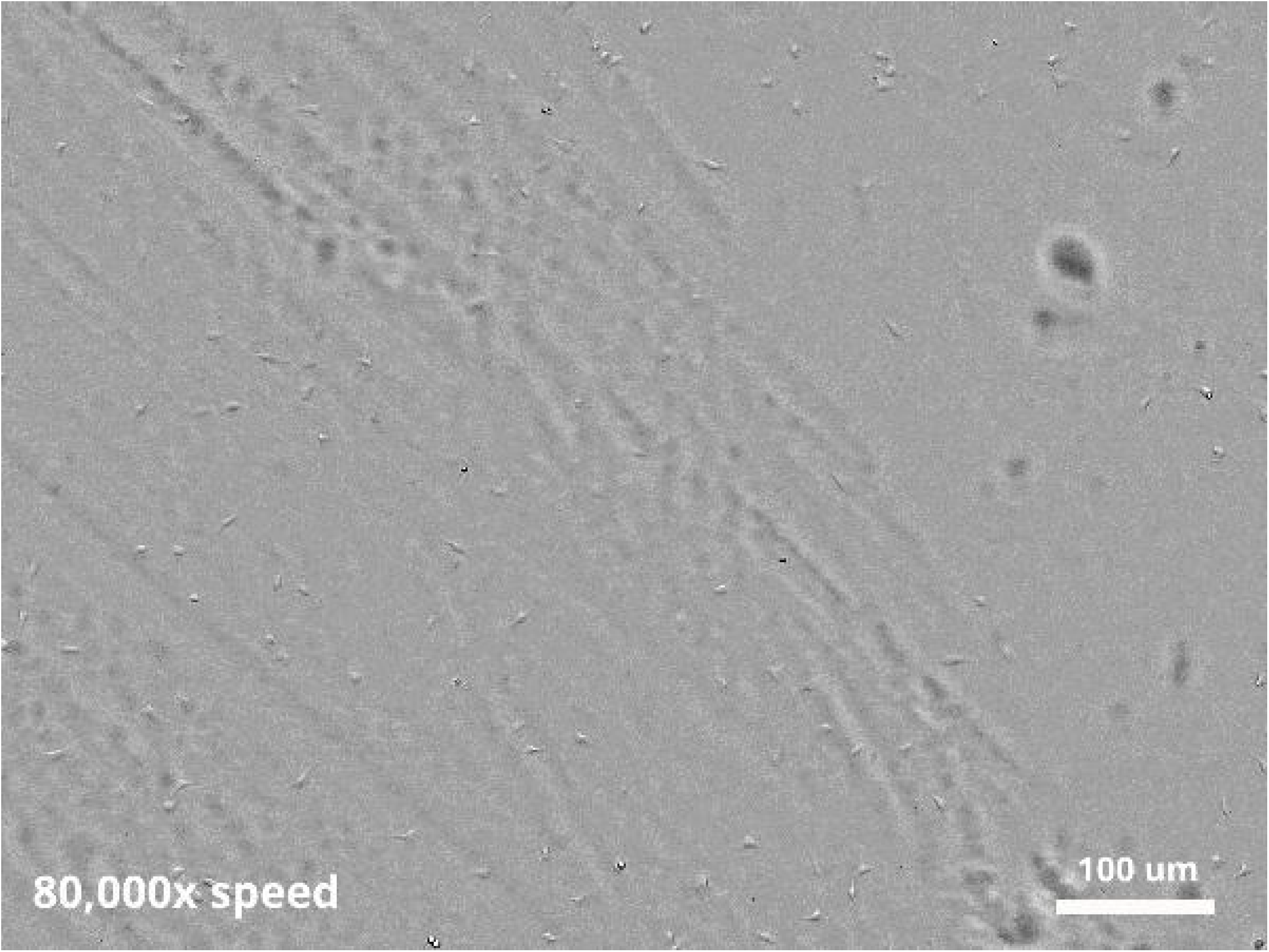

