## additional data and figures for "Simple and effective serum-free medium for sustained expansion of bovine satellite cells for cell cultured meat"

***Supplementary Table 1: B8 formulation and sourcing for this study.***

| Component | Concentration | Supplier | Catalog # |
|---|---|---|---|
| **DMEM/F12** basal media | N/A | ThermoFisher | 11320033 |
| 2-Phospho-**L-ascorbic acid** trisodium salt | 200 µg/mL | Sigma | 49752-10G |
| **Insulin** (human, recombinant) | 20 µg/mL | Sigma | 91077C-250MG |
| **Transferrin** (human, recombinant) | 20 µg/mL | InVitria | 777TRF029 |
| Sodium **selenite** | 20 ng/mL | Sigma | S5261-10G |
| Fibroblast growth factor (**FGF-2**) | 40 ng/mL | PeproTech | 100-18B |
| Neuregulin (**NRG1**) | 0.1 ng/mL | PeproTech | 100-03 |
| Transforming growth factor (**TGFβ3**) | 0.1 ng/mL | R&D Systems | 8420-B3-005/CF |
| UltraPure **Water** | 5.8% (v/v) | ThermoFisher | 10977015 |
| **Antibiotic/Antimycotic** | 1% (v/v) | ThermoFisher | 1540062 |

Functional species in media components are bolded. To prepare media, two aliquots were first prepared. For aliquot 'A,' 400 mg/mL 2-Phospho-L-ascorbic acid trisodium salt was slowly prepared in water, sterilized, and aliquoted in 250 µL portions. For aliquot 'B,' 40 mg/mL insulin was added to water, and 1 N HCl was added until insulin had dissolved. Slowly, 1 N NaOH was added to bring the pH up to ~6. Next, 40 mg/mL transferrin, 40 µg/mL sodium selenite, 80 µg/mL FGF-2, 0.2 µg/mL NRG1, and 0.2µg/mL TGFβ3 were added, and the solution was sterilized and aliquoted in 250 µL portions. To prepare media, DMEM/F12 (500 mL) was sterilized with 5.3 mL of 100x Antibiotic/Antimycotic and 31 mL of water through a sterile filter, and aliquots A and B were added after sterilization. Following protocol previously described for B8 media, media used in this study was used within one month of preparing from frozen aliquots, and media was warmed to room temperature before feeding cells.

*Supplementary Table 2: Summary of all media supplements tested.*

| Component | Supplier | Catalogue # | Range (ng/mL) |
|---|---|---|---|
| Hydrocortisone | Sigma | H0888-1G | 3.125—100 |
| Estradiol | Sigma | E8875-1G | 1—5,000 |
| Progesterone | Sigma | P8783-1G | 0.3125—10 |
| Dexamethasone | Sigma | D4902-100MG | 3.125—400 |
| BMS 564929 | Tocris Bioscience | 5274 | 0.1—500 |
| Bovine growth hormone | MP Biomedical | 02160074.1 | 7.8125—250 |
| Curcumin | Sigma | C7727-500MG | 1—5,000 |
| Spermidine | Sigma | S2626-1G | 100—500,000 |
| Ethanolamine | Acros Organics | 149582500 | 10—50,000 |
| Linoleic acid | Sigma | L5900-10MG | 25—3,200 |
| Oleic acid | Sigma | O1257-10MG | 25—3,200 |
| Stat3 inhibitor | Sigma | 573096-1MG | 100—500,000 |
| Interleukin-6 | BioRad | PBP021 | 0.01—50 |
| Leukemia inhibitory factor | Peprotech | 300-05 | 0.3125—40 |
| Hepatocyte growth factor | Novus Biologicals | NBP2517300.1MG | 0.3125—40 |
| Platelet derived growth factor BB | ThermoFisher | PHG0045 | 0.15625—20 |
| Pigment-epithelium derived factor | R&D Systems | 1177SF025 | 0.390625—50 |
| Insulin-like growth factor | ThermoFisher | PHG0078 | 0.1—50 |
| Cardiotrophin 1 | Novus Biologicals | NBP199745 | 0.15625—20 |
| Recombinant human albumin | Sigma | A9731-1G | 100,000—800,000 |

*Supplementary Figure 1: BSC growth in dilutions of BSC-GM and HiDef-B8*

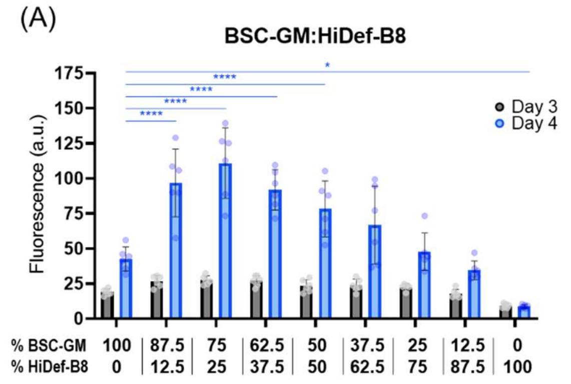


**Figure S1: Short-term growth in BSC-GM mixed with HiDef-B8.** BSC proliferation over 3 & 4 days in mixtures of BSC-GM (20% FBS) HiDef-B8 media. Results show that up to 50% HiDef-B8 significantly improves growth compared to BSC-GM, and that up to 87.5% does not significantly reduce growth compared to BSC-GM. n=6 distinct samples; statistical significance was calculated by one-way ANOVA on day 4 data comparing all samples with BSC-GM controls, and is indicated by asterisks, in which $p < 0.05$ (*), $p < 0.01$ (**), $p < 0.001$ (***), and $p < 0.0001$ (****).

*Supplementary Figure 2: Individual supplement effects on B8 media*

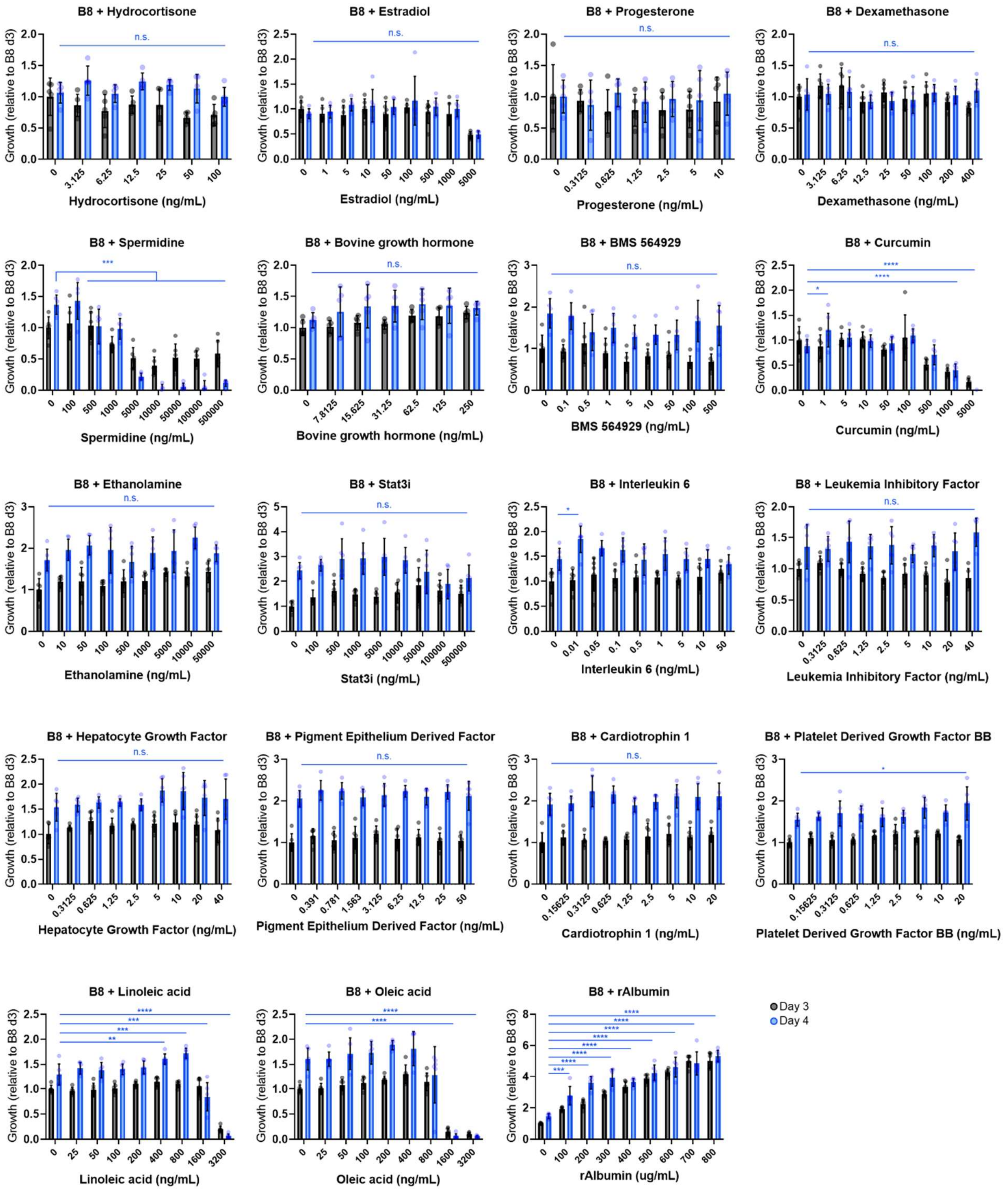


**Figure S2: Short-term growth in supplemented B8 media.** BSC proliferation over 3 & 4 days B8 supplemented with various supplements. Values are fluorescence based and given relative to day 3 B8 values. n=6 distinct samples; statistical significance was calculated by one-way ANOVA on day 4 data with multiple comparisons between supplemented samples and B8 controls, and is indicated by asterisks, in which p < 0.05 (*), p <0.01 (**), p < 0.001 (***), and p < 0.0001 (****).

*Supplementary Figure 3: Combined supplement effects on B8 media*

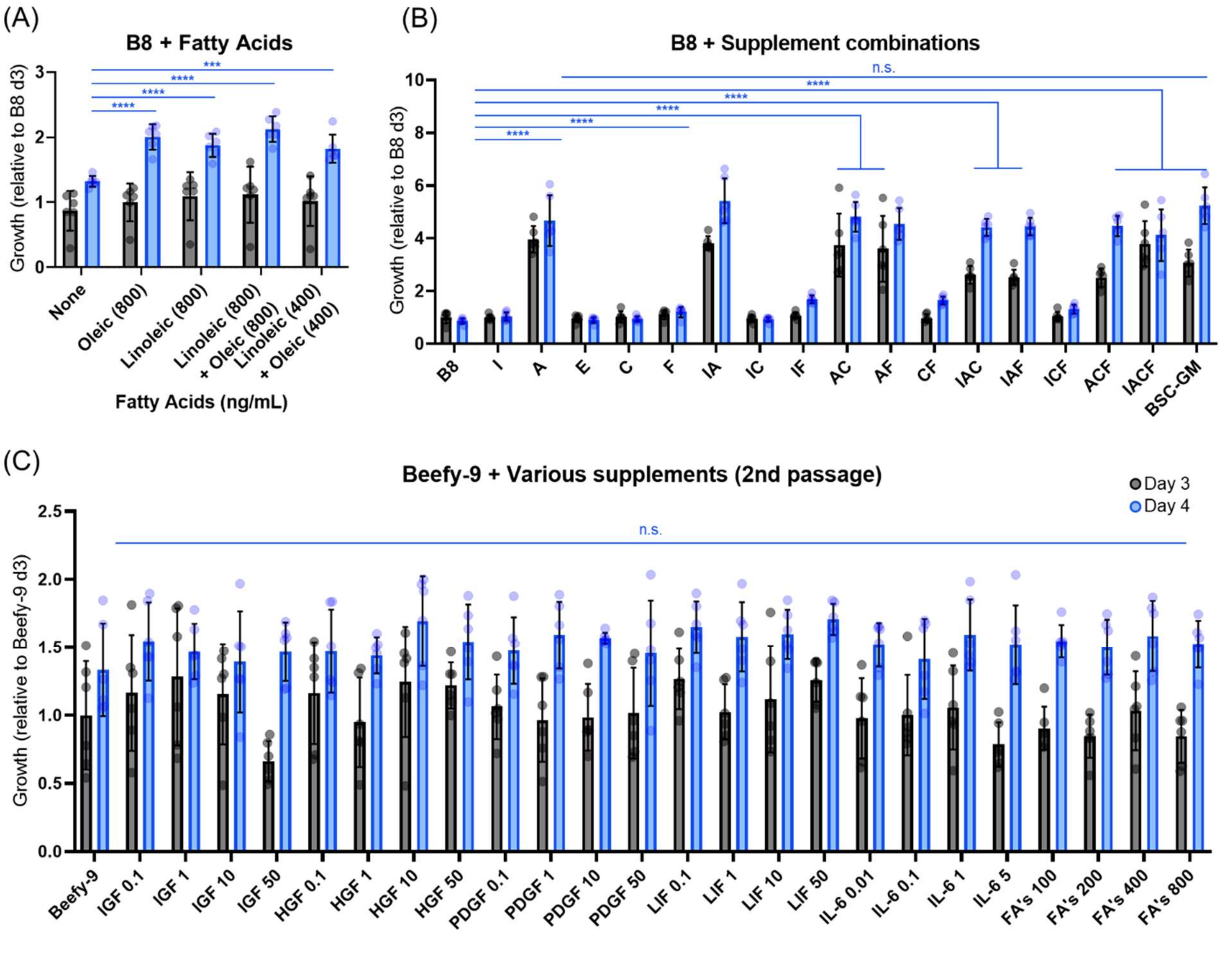


**Figure S3: Short-term growth in B8 media supplemented with combinations of supplements. (A)** BSC proliferation over 3 & 4 days B8 supplemented with combinations of fatty acids. Values are fluorescence based and given relative to day 3 B8 values. n=6 distinct samples; statistical significance was calculated by one-way ANOVA on day 4 data with multiple comparisons between supplemented samples and B8 controls, and is indicated by asterisks, in which $p < 0.05$ (*), $p < 0.01$ (**), $p < 0.001$ (***), and $p < 0.0001$ (****). **(B)** BSC proliferation over3 & 4 days in B8 supplemented with combinations of factors, in which: B8 = B8; I = IL-6 (.01 ng/mL); A = rAlbumin (800 µg/mL); C = Curcumin (1 ng/mL); F= linoleic acid (400 ng/mL) and Oleic acid (400 ng/mL); and BSC-GM = serum-containing growth media. n=6 distinct samples; statistical significance was calculated by one-way ANOVA on day 4 data with multiple comparisons between all samples, and is indicated by asterisks, in which $p < 0.05$ (*), $p < 0.01$ (**), $p < 0.001$ (***), and $p < 0.0001$ (****). Statistically significant differences between B8 and other samples are shown, as is insignificance between rAlbumin supplemented B8 and BSC-GM ($p > 0.9990$). **(C)** BSC proliferation over 3 & 4 days in Beefy-9 supplemented with various factors (IGF = insulin-like growth factor 1 at concentrations from 0.1—50 ng/mL; HGF = hepatocyte growth factor at concentrations from 0.1—50 ng/mL; PDGF = platelet derived growth factor BB at concentrations from 0.1—50 ng/mL; LIF = leukemia inhibitory factor at concentrations from 0.1—50 ng/mL; IL-6 = interleukin 6 at concentrations from 0.01—5 ng/mL, FA's = Combinations of Linoleic & Oleic acid at concentrations ranging from 100-800 ng/mL of each fatty acid). Cells were passaged in B8/Beefy-9 media before testing in order to eliminate the potential for confounding release of adsorbed serum-derived growth factors from tissue culture plastic. n=6 distinct samples; statistical significance was calculated by one-way ANOVA on day 4 data with multiple comparisons between supplemented samples and Beefy-9 controls, and is indicated by asterisks, in which $p < 0.05$ (*), $p < 0.01$ (**), $p < 0.001$ (***), and $p < 0.0001$ (****).

***Supplementary Video 1: Video of cell growth and differentiation***

****See video file in included Data Files****

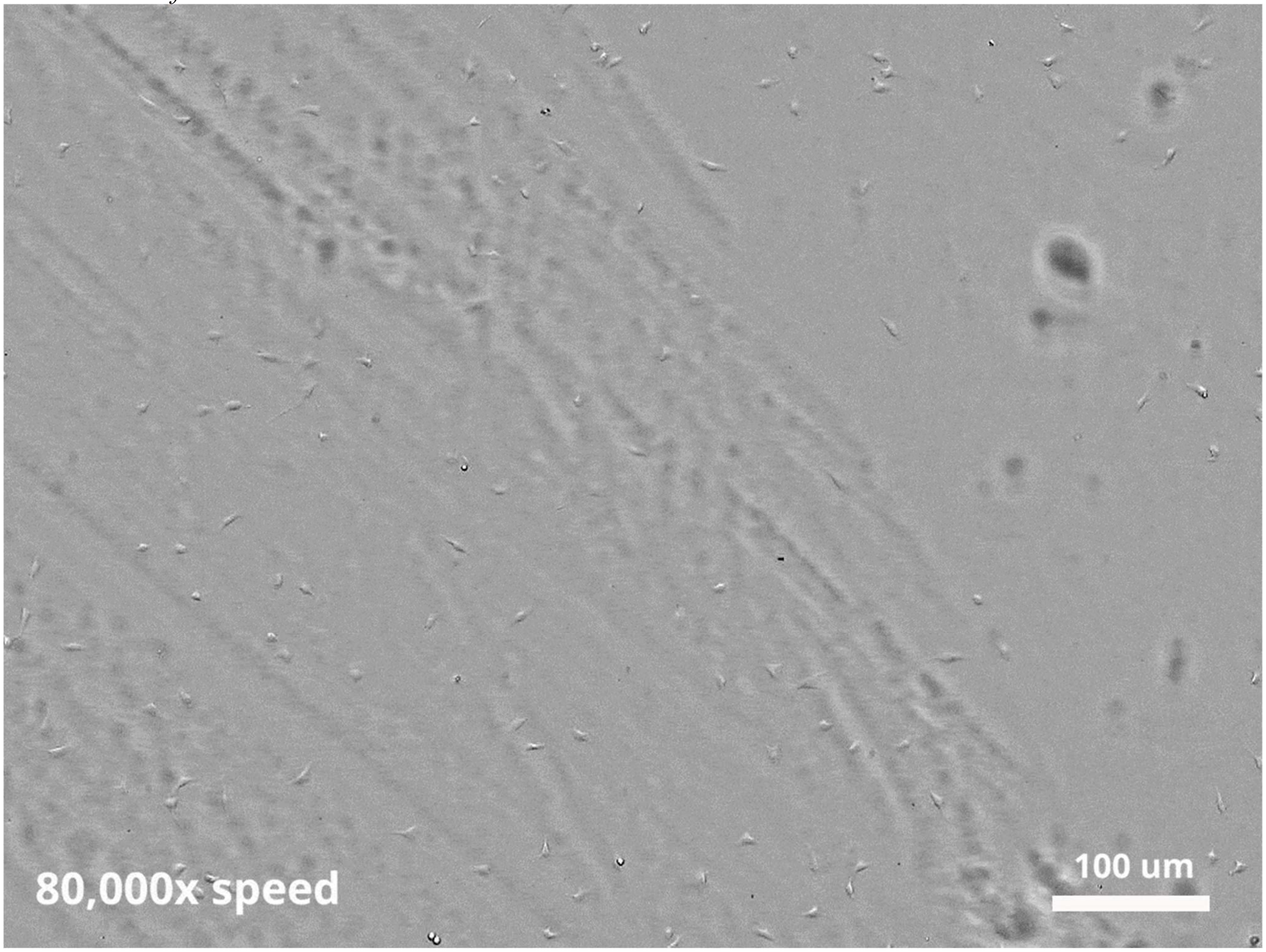


Video of cell growth in Beefy-9 after passaging onto vitronectin-coated plates demonstrates a completely animal-component free culture process for BSCs. Cells were harvested from previous culture vessels with animal-component-free TrypLE, seeding onto new flasks with animal-component-free recombinant vitronectin, proliferated with animal-component-free Beefy-9 media, and differentiated with an animal-component-free differentiation medium. Cells were visualized on an Olympus Provi CM20 over 10 days with imaging every three hours.

*Supplementary Figure 4: Fusion index analysis of differentiated BSCs*

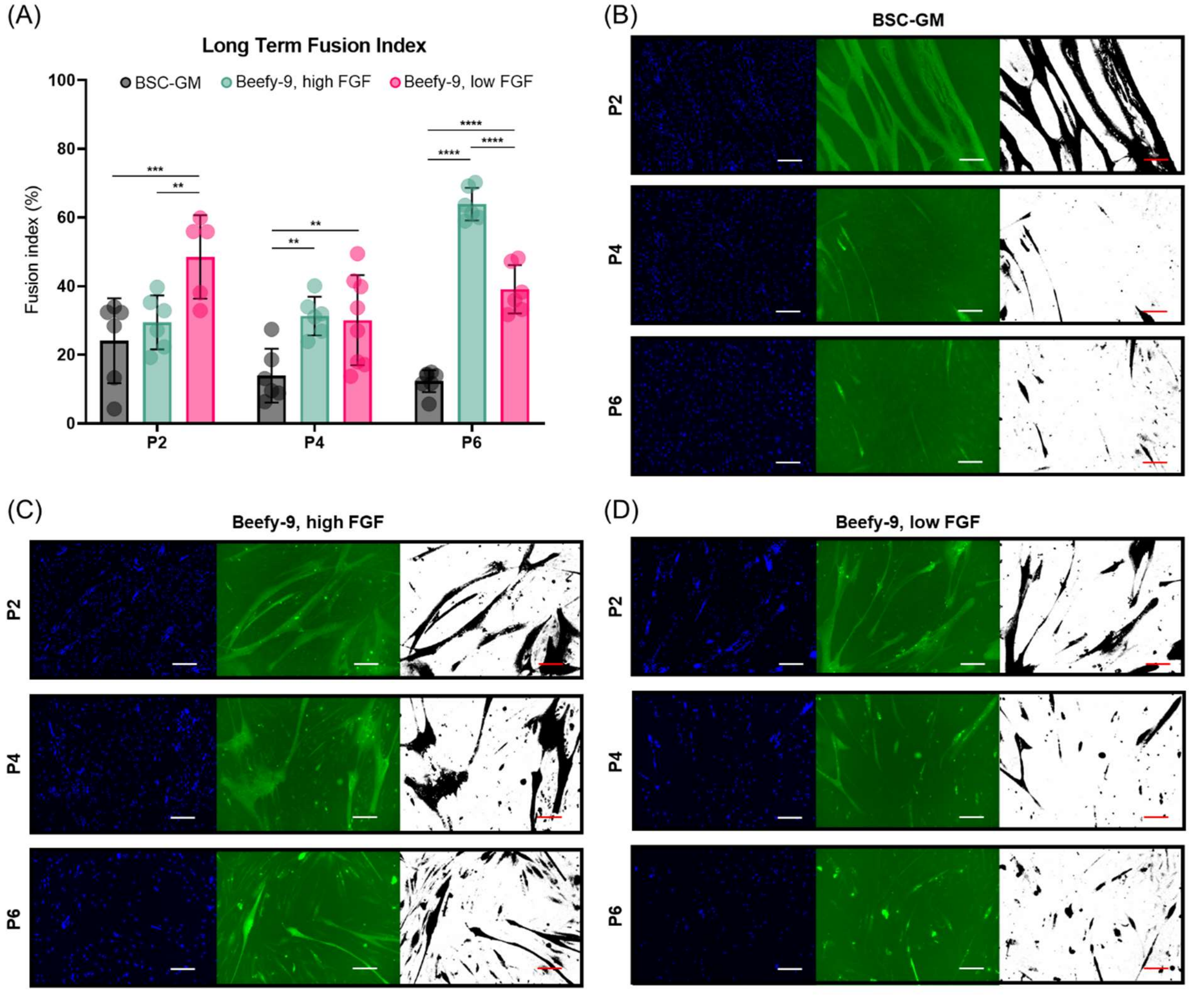


**Figure S4: Fusion index analysis of BSCs cultured over six passages in various media. (A)** Fusion index of images (10x magnification) of BSCs cultured in various media and differentiated for 4 days (P2 and P4) or 6 days (P6). Fusion index given as the percent of imaged nuclei in the images that are superimposed with myosin heavy chain (MHC) staining after thresholding in ImageJ (threshold value = 18 for all images). n=5-8 distinct images depending on whether any images were excluded due to imaging artifacts (e.g., bubbles) that clearly disrupted image analysis; statistical significance was calculated by two-way ANOVA with multiple comparisons between media types for each distinct passage tested, and is indicated by asterisks, in which $p < 0.05$ (*), $p < 0.01$ (**), $p < 0.001$ (***), and $p < 0.0001$ (****). **(B-D)** Representative images (Left = DAPI; Middle = Myosin Heavy Chain (MHC); Right = MHC post-thresholding) of those used for generating figure S4(A) for each media type and passage number tested. Scale bars 100 µm.

### *Supplementary Table 3: Component sourcing and pricing for cost analysis*

| BSC-GM | | | | | | | | | | | | | | | |
|---|---|---|---|---|---|---|---|---|---|---|---|---|---|---|---|
| **This study** | | | | | | | | **Bulk** | | | | | | | |
| Component | Unit / L | Amnt / L | Supplier | Cat. # | Units/order | Cost/order | Cost / L | Component | Unit / L | Amnt / L | Supplier | Cat. # | Units/order | Cost/order | Cost / L |
| DMEM (Liquid)<br>+ HEPES<br>+ Sodium Bicarbonate | mL | 800 | ThermoFisher | 10569044 | 5,000 | 294 | 47.04 | DMEM (Powder) | mL | 800 | ThermoFisher | 12100061 | 10000 | 27.88 | 2.23 |
| | | | | | | | | HEPES | g | 3.574 | Grainger | 30TY36 | 10000 | 1060.8 | 0.38 |
| | | | | | | | | Sodium Bicarbonate | g | 1.2 | Grainger | 31GD57 | 5000 | 120.59 | 0.03 |
| FBS | mL | 200 | ThermoFisher | 26140079 | 500 | 606 | 242.40 | FBS | mL | 200 | Zenbio | SER-500 | 500 | 452 | 180.80 |
| FGF-2 | ug | 1 | Peprotech | 100-18B | 1000 | 800 | 0.80 | FGF2 | ug | 1 | Peprotech | 100-18B | 1000 | 800 | 0.80 |
| **Total** | | | | | | | **290.24** | **Total** | | | | | | | **184.24** |

| Beefy-9 (40 ng/mL FGF-2) | | | | | | | | | | | | | | | |
|---|---|---|---|---|---|---|---|---|---|---|---|---|---|---|---|
| **This study** | | | | | | | | **Bulk** | | | | | | | |
| Component | Unit / L | Amnt / L | Supplier | Cat. # | Units/order | Cost/order | Cost / L | Component | Unit / L | Amnt / L | Supplier | Cat. # | Units/order | Cost/order | Cost / L |
| DMEM (Liquid)<br>+ HEPES<br>+ Sodium Bicarbonate | mL | 1000 | ThermoFisher | 11330057 | 5,000 | 334 | 66.80 | DMEM/F12 (Powder) | mL | 1000 | ThermoFisher | 12500096 | 50000 | 189 | 3.78 |
| | | | | | | | | HEPES | g | 3.574 | Grainger | 30TY36 | 10000 | 1060.8 | 0.38 |
| | | | | | | | | Sodium Bicarbonate | g | 1.2 | Grainger | 31GD57 | 5000 | 120.59 | 0.03 |
| L-ascorbic acid 2-phosphate | mg | 200 | Sigma | 49752-10G | 10000 | 65.5 | 1.31 | L-ascorbic acid 2-phosphate | mg | 200 | Sigma | 49752-100G | 100000 | 417 | 0.83 |
| Insulin | mg | 20 | Sigma | 91077C-250MG | 250 | 254 | 20.32 | Insulin | mg | 20 | InVitria | 777HSA017S-1KG | 100000 | 41920 | 8.38 |
| Transferrin | mg | 20 | inVitria | 777TRF029 | 1000 | 260.5 | 5.21 | Transferrin | mg | 20 | inVitria | 777TRF029 | 10000 | 1823 | 3.65 |
| Sodium selenite | ug | 20 | Sigma | S5261-10G | 10000000 | 32.1 | 0.00 | Sodium selenite | ug | 20 | Sigma | S5261-100G | 100000000 | 32.1 | 0.00 |
| FGF2 | ug | 40 | Peprotech | 100-18B | 1000 | 800 | 32.00 | FGF2 | ug | 40 | Peprotech | 100-18B | 1000 | 800 | 32.00 |
| NRG1 | ng | 100 | Peprotech | 100-03 | 10000 | 80 | 0.80 | NRG1 | ng | 100 | PeproTech | 100-36E | 1000000 | 1500 | 0.15 |
| TGFb3 | ng | 100 | R&D Systems | 8420-B3-005/CF | 5000 | 199 | 3.98 | TGFb3 | ng | 100 | PeproTech | 100-36E | 1000000 | 5200 | 0.52 |
| rAlbumin | mg | 800 | Sigma | A9731-1G | 1000 | 108 | 86.40 | rAlbumin | mg | 800 | ScienCell | OsrHSA | 1000000 | 30703 | 24.56 |
| **Total** | | | | | | | **216.82** | **Total** | | | | | | | **74.28** |

| Beefy-9 (5 ng/mL FGF-2) | | | | | | | | | | | | | | | |
|---|---|---|---|---|---|---|---|---|---|---|---|---|---|---|---|
| **This study** | | | | | | | | **Bulk** | | | | | | | |
| Component | Unit / L | Amnt / L | Supplier | Cat. # | Units/order | Cost/order | Cost / L | Component | Unit / L | Amnt / L | Supplier | Cat. # | Units/order | Cost/order | Cost / L |
| DMEM (Liquid)<br>+ HEPES<br>+ Sodium Bicarbonate | mL | 1000 | ThermoFisher | 11330057 | 5,000 | 334 | 66.80 | DMEM/F12, Powder | mL | 1000 | ThermoFisher | 12500096 | 50000 | 189 | 3.78 |
| | | | | | | | | HEPES | g | 3.574 | Grainger | 30TY36 | 10000 | 1060.8 | 0.38 |
| | | | | | | | | Sodium Bicarbonate | g | 1.2 | Grainger | 31GD57 | 5000 | 120.59 | 0.03 |
| L-ascorbic acid 2-phosphate | mg | 200 | Sigma | 49752-10G | 10000 | 65.5 | 1.31 | L-ascorbic acid 2-phosphate | mg | 200 | Sigma | 49752-100G | 100000 | 417 | 0.83 |
| Insulin | mg | 20 | Sigma | 91077C-250MG | 250 | 254 | 20.32 | Insulin | mg | 20 | InVitria | 777HSA017S-1KG | 100000 | 41920 | 8.38 |
| Transferrin | mg | 20 | inVitria | 777TRF029 | 1000 | 260.5 | 5.21 | Transferrin | mg | 20 | inVitria | 777TRF029 | 10000 | 1823 | 3.65 |
| Sodium selenite | ug | 20 | Sigma | S5261-10G | 10000000 | 32.1 | 0.00 | Sodium selenite | ug | 20 | Sigma | S5261-100G | 100000000 | 32.1 | 0.00 |
| FGF2 | ug | 5 | Peprotech | 100-18B | 1000 | 800 | 4.00 | FGF2 | ug | 5 | Peprotech | 100-18B | 1000 | 800 | 4.00 |
| NRG1 | ng | 100 | Peprotech | 100-03 | 10000 | 80 | 0.80 | NRG1 | ng | 100 | PeproTech | 100-36E | 1000000 | 1500 | 0.15 |
| TGFb3 | ng | 100 | R&D Systems | 8420-B3-005/CF | 5000 | 199 | 3.98 | TGFb3 | ng | 100 | PeproTech | 100-36E | 1000000 | 5200 | 0.52 |
| rAlbumin | mg | 800 | Sigma | A9731-1G | 1000 | 108 | 86.40 | rAlbumin | mg | 800 | ScienCell | OsrHSA | 1000000 | 30703 | 24.56 |
| **Total** | | | | | | | **188.82** | **Total** | | | | | | | **46.28** |
